# Interplay between mechanochemical patterning and glassy dynamics in cellular monolayers

**DOI:** 10.1101/2023.03.24.534111

**Authors:** Daniel Boocock, Tsuyoshi Hirashima, Edouard Hannezo

**Author notes:** Electronic address.

## Abstract

Living tissues are characterized by an intrinsically mechano-chemical interplay of active physical forces and complex biochemical signalling pathways. Either feature alone can give rise to complex emergent phenomena, for example mechanically driven glassy dynamics and rigidity transitions, or chemically driven reaction-diffusion instabilities. An important question is how to quantitatively assess the contribution of these different cues to the large-scale dynamics of biological materials. We address this in MDCK monolayers, considering both mechanochemical feedbacks between ERK signalling activity and cellular density as well as a mechanically active tissue rheology via a self-propelled vertex model. We show that the relative strength of active migration forces to mechanochemical couplings controls a transition from uniform active glass to periodic spatiotemporal waves. We parameterize the model from published experimental datasets on MDCK monolayers, and use it to make new predictions on the correlation functions of cellular dynamics and the dynamics of topological defects associated with the oscillatory phase of cells. Interestingly, MDCK monolayers are best described by an intermediary parameter region in which both mechanochemical couplings and noisy active propulsion have a strong influence on the dynamics. Finally, we study how tissue rheology and ERK waves feedback on one another, and uncover a mechanism via which tissue fluidity can be controlled by mechano-chemical waves both at the local and global levels.

## Introduction

Unraveling the properties of living materials requires an understanding of how active mechanical forces and material properties arise across sub-cellular and tissue scales, together with how these physical properties are integrated with complex biochemical signalling dynamics occurring within and between cells [1–3]. Experiments on *in vitro* monolayers have provided a fertile playground for developing and testing minimal active matter theories of supracellular dynamics, revealing features such as active nematic turbulence [4, 5], glassy dynamics [6–9], unjamming transitions [10–12], as well as mechanical and/or chemical wave propagation [13–24]. So far these different phenomena have been studied largely in isolation from one another, often by employing a numerical framework such as active particle [7] or vertex-based simulation [8]. A core generic feature within these models is the presence of local energy barriers linked to cell re-arrangements (T1 transitions) which can be overcome by active motility forces, giving rise to active glass signatures for the spatio-temporal evolution of cellular velocity/density [9] that have been compared and contrasted to the classical jamming and glass transitions from passive systems.

Another common feature in tissues is the presence of mechanochemical waves, which have been observed across multiple systems [25], for instance as coupled waves of cell velocity, density, mechanical stress as well as activation of the ERK/MAPK signalling pathway *in vitro* in Madin-Darby canine kidney (MDCK) cells [22, 24]. Spatio-temporal ERK waves have also been observed and shown to be critical *in vivo* for bone regeneration in zebrafish [26], murine cochlear morphogenesis [27], and in murine skin wound healing [28]. Recently in [24] we proposed a minimal quantitative one-dimensional model showing how such waves can arise from simple mechanochemical couplings (Fig. 1A,B) via a spatio-temporal instability that can recapitulate a number of mean-field features of wave propagation [22, 24]. However, how to unify such mechanochemical description with previous modelling and experimental observations on the jamming transitions and glassy dynamics seen cell monolayers remains an outstanding challenge.

**FIG. 1:**
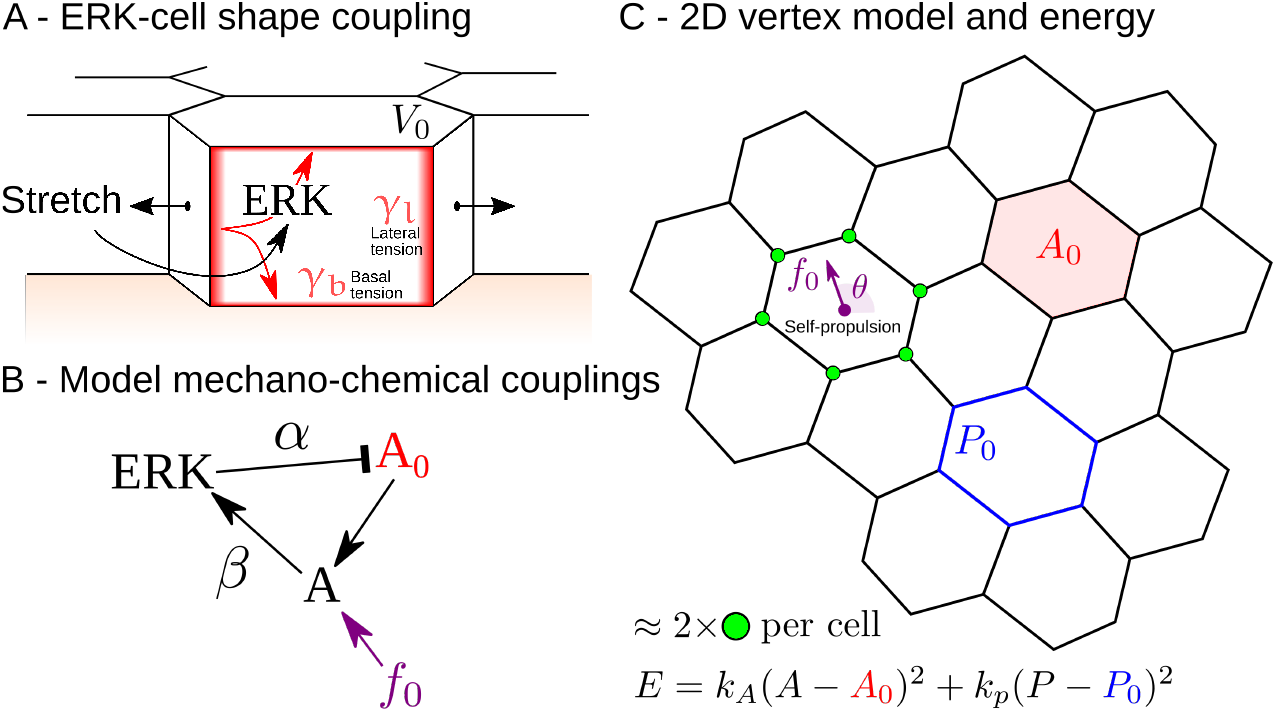
A/ Schematic of the three-dimensional forces exerted on cells in confluent monolayers. In-plane cell area is determined by a balance of cytoskeletal forces on the basal and lateral sides, which can be modulated by ERK signalling activity [22, 24]. B/ Schematic of the mechano-chemical couplings considered in the model. ERK signalling activity impacts on preferred cell area *A*_0_ with coupling strength *β*, creating mechanical stresses for cells with area *A* below or above this preferred value. This, together with stresses from self-propulsion forces *f*_0_ results in changes in area *A* which feeds back on ERK with coupling strength *β*. C/ Schematic of the in-plane 2D vertex model, its associated energy and persistent random motility forces *f*_0_.

In this work, we implement a numerical model for the rheology of active 2D confluent tissues - a self-propelled vertex model - that also incorporates mechanochemical feedbacks between ERK signalling dynamics and cellular density. We show that the relative strength of active migration forces to oscillatory mechanochemical couplings controls a transition from a uniform active glass to one with periodic spatiotemporal patterns. By identifying relationships between mechanochemical coupling parameters and oscillatory amplitudes of ERK and density, we are able to parameterize the model from published experimental datasets on MDCK monolayers *in vitro* [22, 24]. Then we show that a number of quantitative features in the datasets, including temporal auto-correlation functions and the dynamics of topological defects associated with the mechanochemical phase of cells, are recapitulated in an intermediary parameter region where both ERK-density oscillations and noisy active propulsion play an important role. Finally, we study computationally how T1 transition frequency, and thus local tissue fluidity, is subject to spatio-temporal control by oscillatory mechano-chemical activity, and discuss the relevance of this model to other biological settings.

## Results

### Active vertex model with mechanochemical feedbacks

We start by considering the typical vertex model energy, which provides a minimal description for a confluent epithelial monolayer by assuming that 2D cell shapes arise from a balance between adhesive and tensile forces at the cell-cell contacts [10, 29–31]:

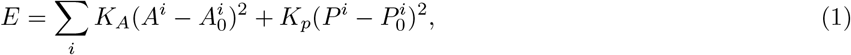

where the first term models a target cellular area 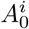while the second term assumes that cells have a preferred perimeter 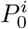based on a balance between cell-cell adhesion and cytoskeletal driven tension (Fig. 1C). A more mathematical viewpoint sees this preferred perimeter as the first non-linear stabilizing terms allowed by symmetry. An important result concerns the value of the shape index 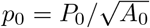 which in homogeneous systems controls the tissue fluidity [10]: systems with *p*_0_ ≲ 3.81 are in a solid state characterized by energy barriers for cell re-arrangements, systems with *p*_0_ ≳ 3.81 are in a fluid state where rigidity is lost. To implement the dynamics of this model numerically, we used the cell-based Chaste library [32] to develop a mechanochemically coupled vertex model. Motion of tri-cellular junctions 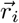 was described by overdamped dynamics, 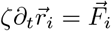 where the time-scale of stress propagation is affected by friction coefficient ζ with the substrate, and the force 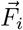 derives from the vertex model energy described above.

Although investigations of the vertex model have concentrated mostly on monolayers constituting cells with constant mechanical properties and self-propulsion, some works have started to investigate the effect of additional variables such as biochemical signalling, morphogen diffusion or mechanosensation [33–35]. Here, we incorporate and study the effect of mesoscopic oscillatory dynamics using mechanochemical ERK waves as a basis. To do so we generalize our previous one-dimensional mechanochemical model [24] to 2D cells with target areas 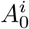 dependent on local ERK signaling activity *E*_*i*_ with delay *τ*_*A*_ (Fig. 1A,B):

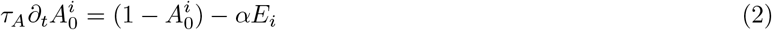

where length scales have been non-dimensionalized using the average cell area and ERK activity non-dimensionalized and centered by its steady-state activation. α is a coupling strength that controls how much ERK signalling impacts on the preferred cell area A_0_ (i.e. signalling to mechanics), and has been shown to be positive from optogenetic ERK activation experiments [24].

An additional equation on ERK activity is required to close to the system, which based on previous experimental findings [22, 24], we take as proportional to local area *A*_*i*_ with a delay *τ*_*E*_:

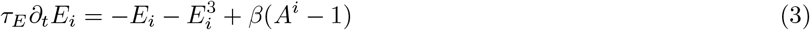

where *β* is a coupling strength that controls the sensitivity of ERK activation as a function of cell area (i.e. mechanics to signalling), and has been shown to also be positive from monolayer stretching experiments [22]. This 2D model is identical at linear order to the one that we studied analytically in 1D in [24], and leads to spatio-temporal instabilities for a critical value of the product of mechanochemical couplings (*αβ*)_*c*_. Physically, this arises due to an oscillatory instability in a feedback loop in which changes in actual area *A*^*i*^ are mechano-sensed via ERK signalling to modify target areas 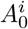, with energy minimization closing 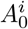back onto *A*^*i*^. To constrain parameters, we take the time scales τ_*A*_ and τ_*E*_ as previously inferred [24], the ratio *K*_*A*_/K_*P*_ from the literature [9, 36] and left the ratio of elasticity to substrate friction K_*A,P*_ /ζ as a free parameter in order to match the wavelength in data (*∼* 20 cells - see Table S1). Running simulations for different values of the coupling constants α and β confirmed that regular spatio-temporal waves emerge above a critical value of αβ which depended on the shape index *p*_0_ (Fig. S1A,B), and we also found that different ratios of α/β dictate the relative amplitudes of mechanical over ERK activity waves (Fig. 2A,B).

**FIG. 2:**
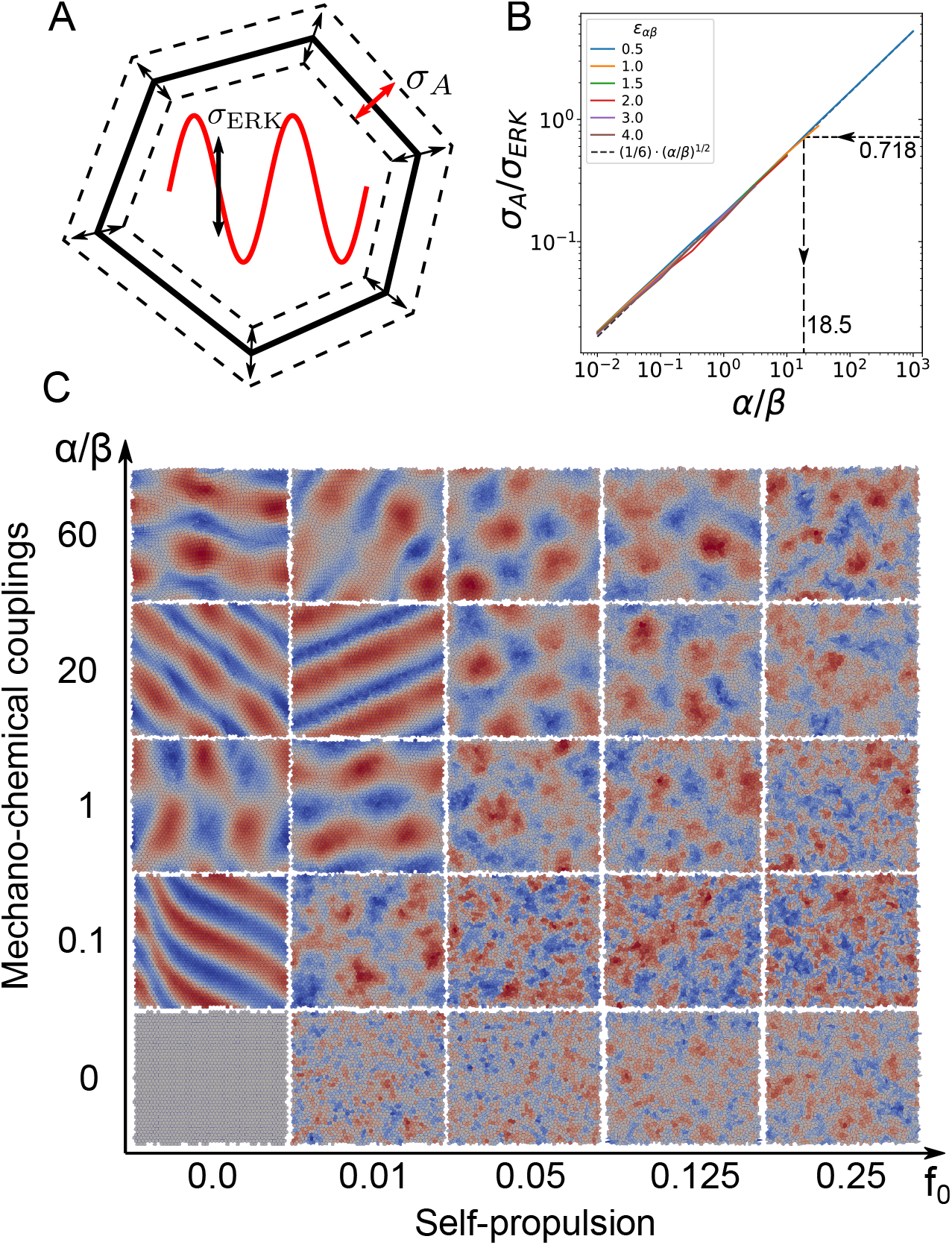
A,B/ Relative amplitude of area to ERK oscillation *σ*_*A*_*/σ*_*ERK*_ vs ratio of mechano-chemical coupling constants *α/β*, showing that different values of the product *αβ* robustly scale along the same master curve, allowing us to fit *α/β* from the experimentally observed value of *σ*_*A*_*/σ*_*ERK*_ (dashed lines, see also Fig. S3B). C/ Phase diagram of mechano-chemical patterning in a 2D vertex model with ERK signalling activity and self-propulsion, as a function of active self-propulsion forces *f*_0_ (which introduce persistent noise in the system) and the relative strength of mechanical and chemical couplings *α/β* (see Fig. S4 for a phase diagram as a function of the product *αβ*).

To compare features more quantitatively with data, we sought to use these findings to better constrain parameters α and β. To constrain the ratio α/β, we reasoned that we could use the relative amplitudes of area and ERK signalling activity oscillations (resp. σ_*A*_ and σ_*E*_) in our datasets. Interestingly, plotting this ratio for different values of α and β revealed that σ_*A*_/σ_*E*_ scaled with the ratio of α/β, independent of the value of αβ, so that all tested values of αβ could be rescaled on the power law σ_*A*_/σ_*E*_ *∝* (α/β)^1*/*2^ (Fig. 2B). This provides a robust method for constraining α/β, as σ_*A*_/σ_*E*_ was highly consistent over the three repeats of previously published datasets on mechanochemical ERK patterning in confluent M DCK monolayers [24] (Fig. S 3B), allowing us to estimate α/β *≈* 18.5 (Fig. 2B).

Despite such parameterization being able to match several qualitative features and summary statistics of the data (Fig. S1C and Fig. S2B-C), visual comparisons to experimental datasets (Fig. 2C) make it clear that ERK waves in MDCK monolayers are much more disordered than the ones we expect for this minimal model in the noise-free limit. Although a number of sources of noise could be considered - ranging from noise in ERK signalling to junctional fluctuations - previous modelling [9] h as shown that a number off eatures of MDCK collective dynamics can be explained by a model of glassy dynamics driven by noisy active migration forces. Thus, in addition to the mechanochemical model above, we considered active cell migration in direction 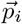 (unit vector) with characteristic force *f*_0_ exerted on the substrate [8, 37]. Writing force balance on cell centers 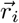gives

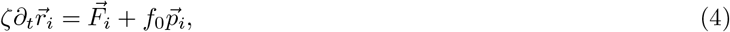

where the angle of polarization *θ*_*i*_ of the polarity unit vector 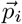 evolves with zero mean Gaussian white noise as ∂_*t*_θ_*i*_ = η_*i*_(t) giving a cellular persistence time 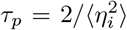which we set as τ_*p*_ = 30min based on published data on the time scale of traction force changes [22]. In the vertex model implementation, the propulsion force for each cell is applied to all of the vertices defining its tri-cellular junctions. In practice the self-propulsion force always appears with substrate friction as the ratio f_0_/ζ which we simply refer to as f_0_ in the rest of this text.

### Effect of self-propulsion and mechanochemical couplings on monolayer dynamics

In the absence of mechanochemical couplings *α* = 0 (where we keep *β ≠* 0 so that ERK signalling is a passive variable that tracks cell area), we find, a s expected from classical work [8], that a solid monolayer *p*_0_ ≲ 3.81 can be fluidized by active traction forces above a critical value f_0_ (Fig. S1D and Movie S1), leading to collective streaming and locally disordered cellular shapes characteristic of glassy dynamics [8, 9]. However, in this *α* = 0 case, ERK signalling does not show any spatial or temporal periodicity (Fig. S2A, see also [36]), irrespective of whether f_0_ is above or below the unjamming threshold, which contrasts with experimental observations of ERK patterning [22, 24]. Thus, the experimental dynamics did not fully match the cases of active jamming or oscillatory mechanochemical signalling alone, and we hypothesised that the system might be better described by a combination of the two phenomena.

To test this, we first explored the phase diagram of possible patterns in ERK signalling and cell area for different values of *α, β* and *f*_0_ (Fig. S4). As expected, stronger mechanochemical couplings *αβ* favored periodic instabilities while large random migration forces *f*_0_ favored noisy glass-like dynamics with unstructured ERK patterns. Furthermore, we also found that larger ratios *α/β* favoured periodic patterning (Fig. 2C): this is because larger *α/β* increases the relative amplitude of area to ERK oscillations, which makes the system more resistant to area perturbations induced by migration forces.

We then repeated our analysis of ERK and area amplitudes for fitting mechano-chemical couplings *α* and *β* in the presence of self-propulsion noise f_0_, finding that our previous estimate for *α/β* from amplitude ratio *σ*_*A*_/*σ*_*E*_ was quite robust to different values of *f*_0_ (Fig. S3C,F). Furthermore, by jointly matching amplitudes from *σ*_*A*_, *σ*_*E*_ and *σ*_*A*_/*σ*_*E*_, we could estimate a region of parameter space, *ϵ*_*αβ*_ = (*αβ − αβ*_c_)/*αβ*_c_ *≈* 0.8 and *f*_0_ = 0.125 *−* 0.15, where our model was fully parameterized from data (Fig. S3D-F and see simulation in Movie S1). This quantitative analysis of amplitudes, along with qualitative inspection of simulations, suggested that intermediate values of α/β and f_0_ could match experimental datasets.

### Quantitative comparisons between simulations and MDCK experiments

To confirm this, we sought to make further quantitative spatio-temporal analyses of the mechanochemical patterns formed across different conditions. Inspired by a previous analysis of subcellular biochemical waves in starfish egg cells [38], we took advantage of the periodicity of ERK signalling to define a phase ϕ for the oscillation in every cell (Fig. S7), which could be mapped spatially and temporally across the entire tissue (Fig. 3A-C). We then studied the effect of different types of activity on the patterns by tracking the motion, and creation-annihilation dynamics, of *±*1 vortex defects corresponding to singularities in the phase (see Movies S2-4), which provide a simpler metric to characterize the complex two-dimensional patterns of ERK activity.

**FIG. 3:**
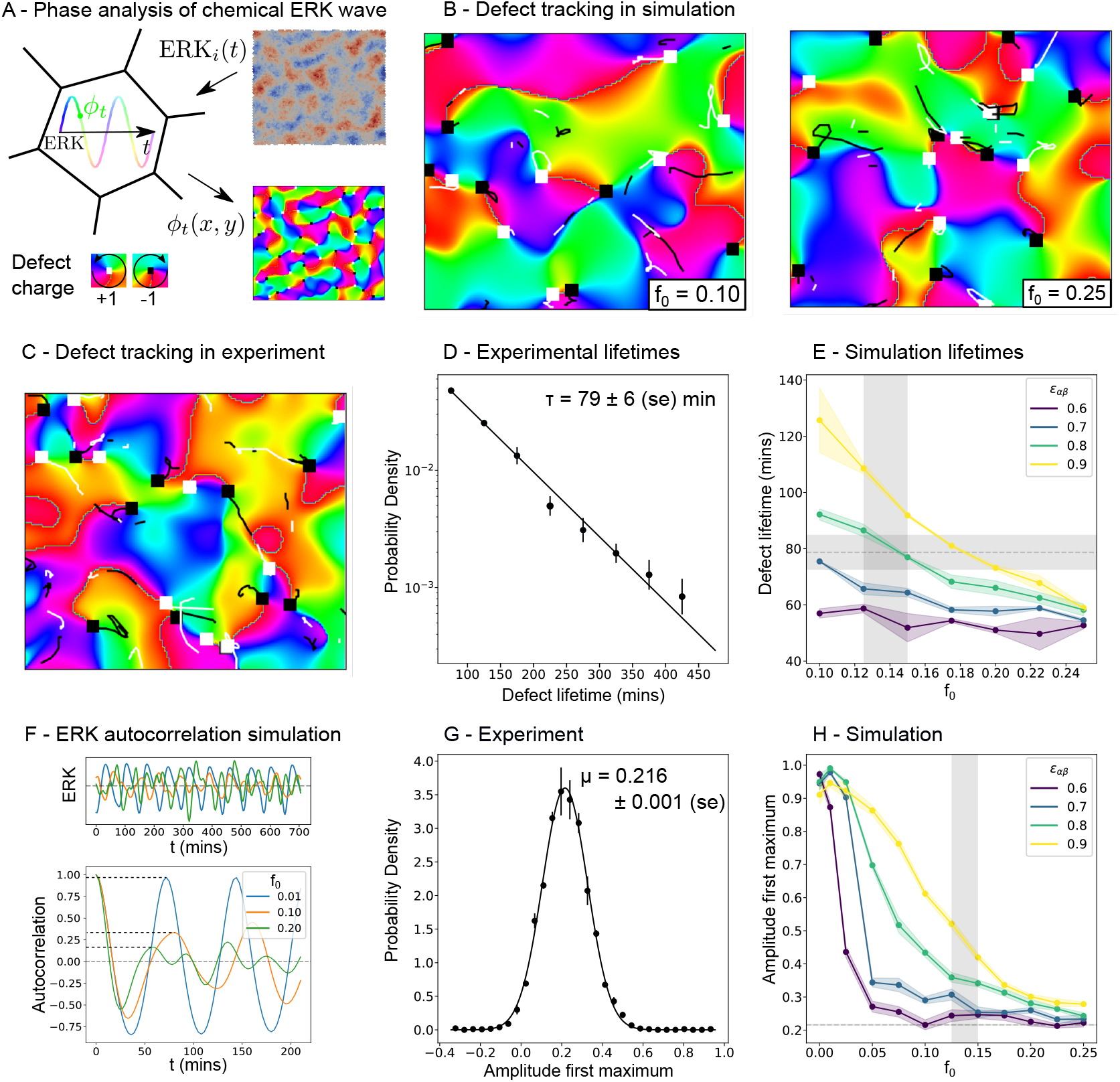
A/ Overview of the tissue-level analysis of ERK patterning. Local cellular phases of ERK signalling activity are smoothed into continuous fields, which have associated topological defect with opposite charge associated to them. B/ Phase-field of ERK activity (color-coded for phase), for different values of active self-propulsion *f*_0_ together with the identification and tracking in time of topological defects (white and black squares indicate resp. +-1 charge.) C/ Phase-field of ERK activity in MDCK monolayers, with the same analysis as in B. D/ Experimental life-time of ERK phase topological defects in MDCK monolayers, showing a good fit to an exponential distribution as observed in simulations. E/ Average life-time of ERK phase topological defects in the simulations as a function of active self-propulsion *f*_0_, for different values of mechanochemical coupling strength *ϵ*_*αβ*_ = (*αβ* − *αβ*_c_)/*αβ*_c_. Shaded intervals indicate experimental values of mean defect lifetime and estimates of *f*_0_, showing that model values of *ϵ*_*αβ*_ *≈* 0.8 and *f*_0_ = 0.125 *−* 1.5 explains the experimental lifetime data while also being consistent with predictions of *α, β* and *f*_0_ from ERK and area amplitudes (Fig. S3D-F). F/ Representative tracks of ERK dynamics in the simulations for different values of *f*_0_ (top) together with corresponding auto-correlation functions (bottom), showing that larger active self-propulsion noise (larger *f*_0_) causes rapid decays, as quantified by the amplitude of the first peak (dashed horizontal lines). G/ Distribution of first-peak amplitudes of ERK autocorrelation across MDCK cells (see Supplementary Information for details). H/ First-peak amplitude of the ERK autocorrelation in simulations as a function of active self-propulsion *f*_0_ and mechanochemical couplings *ϵ*_*αβ*_. Shaded intervals indicate estimates of *f*_0_ as in E and the experimental measurement of first-peak amplitude from G.

We found that the dynamics of the topological defects was markedly different for different values of our parameters. While for all parameter values, the distribution of defect lifetimes was well fitted by an exponential (as in [38]), the average lifetime was strongly dependent on both the mechanochemical coupling strengths *α* and *β* and migration force *f*_0_ (Fig. 3E and S5A) with higher *f*_0_ resulting in higher effective diffusion of topological defects and thus higher probability of defect annihilation. On the other hand, higher overall mechanochemical coupling strength αβ (Fig. 3E) or coupling ratio *α*/*β* (Fig. S5A) favoured regular oscillations which translate into more regular motion of defects and longer lifetimes. Strikingly, going back to the experimental data, we found that the predicted exponential scaling provided a good fit for the experimental defect life time distribution (Fig. 3D). Furthermore, by confronting the predicted and experimentally observed average lifetimes, we found good agreement, thus independently validating our estimates of *f*_0_, α and β (Fig. 3E). This provides additional evidence that the data is consistent with a mixed regime of active glassy dynamics and mechanochemical patterning.

Next, we reasoned that the decay in the temporal auto-correlation of ERK signalling could be used as an additional test of the model. Indeed, as expected from visual inspection of our simulations (Fig. 2C - first column), the noise-free limit displayed highly periodic autocorrelation functions with little decay (Fig. S2B), which could be quantified by the amplitude of the first maximum (Fig. 3F-H). Increasing migration noise *f*_0_ resulted in a gradual loss of periodicity, with a related decay in the amplitude of the first peak. When comparing to data, self-propulsion values in the range *f*_0_ = 0.2 *−* 0.25 could explain the experimental correlation, a somewhat higher value than inferred from other metrics, which could be due to simulations only considering a single source of noise (persistent random migration), whereas others such as cell-cell heterogeneity and cell divisions could be present experimentally.

### Mechanochemical waves modulate the global and local fluidity of the tissue

Finally, we wished to understand theoretically how the solid or fluid state of the monolayer affects mechanochemical waves in the absence of self-propulsion, i.e. for *f*_0_ = 0. We simulated over a range of *αβ* for different values of the preferred cell perimeter *p*_0_ = 3.5 *−* 3.9, finding that solidification leads to a later onset of instability but otherwise qualitatively the same collective mechanochemical oscillations (Fig. S1A-C). This implies that ERK mechanochemical patterning is fairly robust to the rheological state of monolayers, which is likely due to the fact that ERK is only coupled to the local density of cells, something strongly constrained by the confluent assumption of the vertex model.

However, we reasoned that changes in cellular areas and velocities created by mechanochemical waves might feedback on the fluid/solid-like material properties of the monolayer, for instance by locally changing or overcoming the energy barriers related to topological T1 re-arrangements. To test this, we went back to the solid regime *p*_0_ = 3.5 *−* 3.8, which is fluidized by critical values of *f*_0_ as previously described [8]. We then examined whether values of mechanochemical couplings *αβ* might modify the threshold of fluidization/unjamming. We found that large values of *αβ* decreased the threshold of motility force *f*_0_ required for unjamming, consistent with oscillations providing an additional source of effective noise, and that this effect was amplified for the lower value of *p*_0_ = 3.5 - Fig. S6A,B.

Furthermore, we found that the timing and location of T1 transitions was strongly affected by the local phase of mechanochemical waves. By calculating the probability of T1 transitions as a function of the ERK phase ϕ, we indeed observed a two- to three-fold change in the frequency of topological re-arrangements between low and high ERK states (Fig. 4B), which was robust to changes in migration forces *f*_0_. This can be rationalized by considering that area and ERK are closely temporally correlated (ERK lagging slightly behind area with time scale τ_*E*_), so that low-ERK states are also low-area states with low target area and correspondingly high shape index 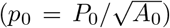. Thus, these regions have higher geometric incompatibility (or frustration) [10] and a propensity towards T1 transitions. The result is that ERK waves can drive local cycles of fluidization and solidification in the monolayer with material properties controlled via patterns of signalling.

**FIG. 4:**
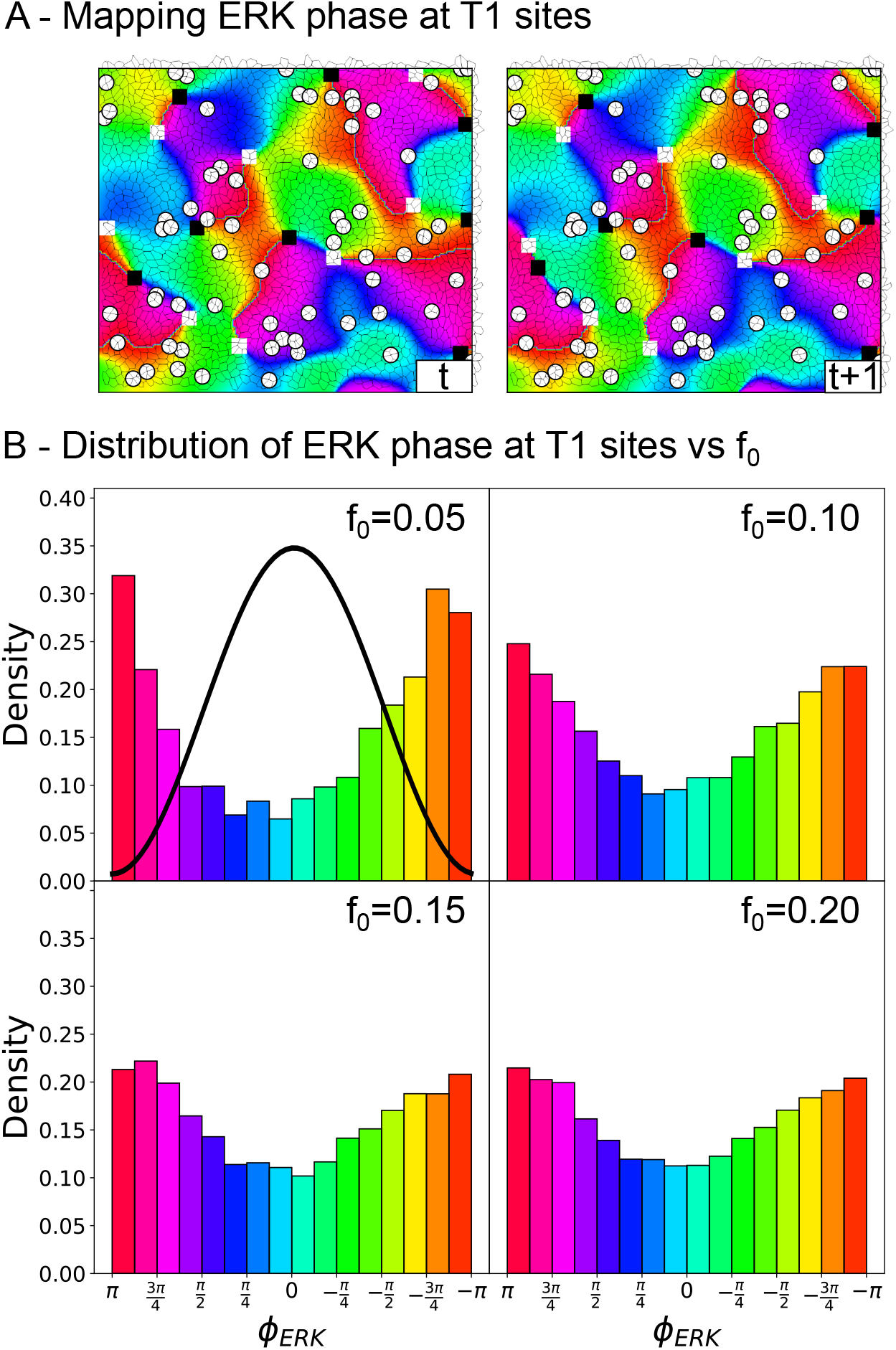
A/ Representative snapshots for the mapping of cell-cell rearrangements (T1 transitions - red circles) to local ERK activity (or phase, see Fig. S7 for color-code) in vertex-model simulations. B/ Distribution of T1 transition probability as a function of ERK phase, for different values of active self-propulsion force *f*_0_, showing strong anti-correlation with ERK activity (black curve shows absolute value and color shows phase) and T1 transitions.

## Discussion

In this work, we have explored the interplay between tissue mechanics and biochemical signalling in a minimal two-dimensional vertex model of a confluent monolayer. Vertex models have been extensively used in the past few years as minimal descriptions of 2D and 3D tissue organization, identifying in particular key geometrical signatures of rigidity transitions, both in the absence of noise and with active self-propulsion forces [8, 10, 29, 39, 40]. In the latter case, it makes specific prediction on glassy dynamics which have common and divergent features compared to the classical glass transition. Interestingly, a number of signatures of active glassy dynamics appear conserved across different modelling frameworks (e.g. vertex vs particle-based models) [7–9], however, these models have historically concentrated on either tissues formed from a single cell type [10, 39], or on the demixing of two cell types possessing constant and defined mechanical properties [41–43].

Here we have considered a model which combines a glassy rheology with an emergent patterning phenomenon that specifies spatio-temporally varying mechanical properties across a tissue. More specifically, we have considered cells with both active self-propulsion and ERK-density mechano-chemical couplings, by associating to each cell a temporally fluctuating signalling activity, and bidirectionally coupling this activity to local cellular mechanics. We concentrate on ERK/MAPK signalling, as this is a key pathway for force-sensing, force-generation and a number of key cellular processes [44–52] (e.g. migration, contractility, differentiation and apoptosis) and because previous works in MDCK monolayers have dissected the types of mechano-chemical couplings that give rise to waves of ERK activity and cell density [22, 24]. However, how to characterize in two dimensions the complex spatio-temporal patterns observed experimentally was not clear. The simulation framework that we propose here, and the associated quantitative analysis based on oscillatory amplitudes, topological defects and auto-correlation functions has allowed us to characterize this system and is highly general to any pathway which forms spatio-temporal patterns by interacting with cellular mechanics, for instance YAP/TAZ [18]. From a more theoretical point of view, this provides an example of pulsating active matter, which has been recently explored via particle-based simulations with oscillating radii and continuum theory [53]. Systematically comparing predictions of our findings to different implementations of tissues, such as particle-based models [53] or phase-field models [54], with distinct rheological properties would be an interesting next step.

Exploring the phase space of possible patterning instabilities revealed a number of interesting features. While the relative strength of biochemical-to-mechanical vs mechanical-to-biochemical coupling (*α*/*β*) plays a key role in determining the relative amplitude of ERK vs cell density oscillations, the ratio of self-propulsion forces *f*_0_ to both global mechano-chemical coupling strength *αβ* as well as to mechano-chemical coupling ratio *α*/*β* determines whether the system is closer to a uniform active glass or a periodic patterning state. Interestingly, we find that multiple signatures of the data - including amplitudes of biochemical and mechanical oscillations, the dynamics of topological defects in the phase of ERK signalling and the decay in autocorrelation of ERK signalling - are consistent with MDCK monolayers being in a intermediate regime characterized by a combination of ERK-density waves and noisy active migration, providing a potentially unifying framework to previous modelling of these types of dynamics [6, 9, 24].

In the future, it will be interesting to test the role of additional sources of fluctuations in addition to cell migration, for instance in ERK signalling dynamics itself or on cell mechanics (i.e. shape index, junctional tensions [55], or average cell density), as well as the effect of time-independent cell heterogeneity (i.e. in active migration force [56] or in proliferation-dependent cell size [57]). Other interesting directions would be to incorporate the effect of active nematics and turbulence [58–62] or to consider alternative theoretical modes of mechano-sensitive couplings which have been recently proposed, for instance on junctional tensions driving active cell rearrangements [34] or junction remodelling [34, 63, 64]. Finally, while we have considered a minimal model of ERK dynamics and its coupling to mechanics, considering more complex features of the pathway such as activator-inhibitor dynamics [65] would be an important next step.

## Materials and Methods

Numerical simulations of the model were performed using the CHASTE library [32]. Analysis was done using custom-codes in Python. Additional details on the numerical procedure, parameter-fitting and quantification strategy can be found in Supplementary Information. All code and data are available upon request.

## Supporting information

Supplementary Theory Note

## Acknowledgements

We thank all members of the Hannezo group for discussions and suggestions. This work received funding from the European Research Council (ERC) under the EU Horizon 2020 research and Innovation Programme Grant Agreement no. 851288 (to E.H.).

## Author contributions

Conceptualization: DB, EH. Simulations: DB. Analysis: DB, TH. Manuscript writing: EH, DB. Manuscript editing: all authors.

