## Supplementary Theory Note for "Interplay between mechanochemical patterning and glassy dynamics in cellular monolayers"

### Materials

Experimental data on cellular tracking and ERK activity measurements in confluent MDCK monolayers can be found in a previous publication [1] with experimental methods described therein. Briefly, the data consisted of three repeats consisting of roughly 3,000 cells each with imaging performed at 5 minute intervals over total times of at least 19 hours. Fluorescent markers captured ERK signal in each cell. Cell center tracking and calculation of Voronoi areas was performed using Fiji plug-in TrackMate [2].

### Methods

#### Vertex model simulations

We used the cell-based Chaste library [3] to develop a vertex model simulation implementing ERK-mechanochemical feedbacks and random persistent migratory forces described in the main text (Eqs. 1-4 and see Fig. 1B,C). Each cell was assigned variables for ERK activity, rest area $A_0$ and self-propulsion angle $\theta$. Vertices were initialized in a hexagonal lattice of 44x44 cells with periodic boundary conditions and initial values ERK = 0 and $A = A_0 = 1$ with length scales normalized by the average cell area ($L = \hat{A}^{1/2}$). Initial values of $\theta$ were chosen uniformly at randomly for each cell.

The following initial Gaussian noise conditions were used so that we could observe instability across all parameter conditions (important when $f_0 = 0$ but carried throughout for consistency). The ratio $\alpha/\beta$ controls the sensitivity of the system to perturbations in area and ERK so to keep initial perturbations small we scaled as follows: for $\alpha > \beta$ we applied the initial perturbation to $ERK$ from the distribution $N(0, 10^{-4}/\alpha^2)$; for $\beta > \alpha$ we applied the initial perturbation to vertex positions from the distribution $N(0, 10^{-4}/\beta^2)$. In both cases we also applied a much smaller amount of initial noise to the less sensitive variable.

Simulations were run with a time-step one hundred times smaller than the smallest time-scale in our model ($dt = 0.01\times\tau_E$ min) for both the vertex step and Euler steps on ODEs for ERK and $A_0$ as well as the SDE for $\theta$. Unless stated otherwise, simulations used a burn-in time of 100 times the predicted period of oscillation ($P = 71$ min, determined from linear stability by $\tau_A$ and $\tau_E$ - see [1]). Data was then recorded over a window of 10 periods, sampling every 50

time steps (every 3 minutes). Longer simulations were required in order to track long-lived topological defects and collect sufficient statistics to plot lifetime distributions (Fig. 3E and S5A). For this we used a burn-in time of 200 periods and recording window of 500 periods. Longer simulations were also required to plot the slower decay in ERK auto-correlation for $\alpha = 0$ (Fig S2A), where we used burn-in time of 100 periods and recording window of 50 periods.

For non-zero $\alpha$, cells possess variable rest areas ($A_0 = A_0(x, t)$ across the tissue), however average and equilibrium rest areas remain close to unity (see Eq. 2) allowing us to control how we vary the shape index $p_0 = P_0/\sqrt{A_0}$ across different simulations through the preferred perimeter $P_0$. Unless otherwise stated simulations used $P_0 = 3.8$ placing them close to the glass transition [4]. We also varied parameters $\alpha$, $\beta$ and $f_0$ across our simulations while parameters summarized in Table S1 were held fixed.

### Amplitude analysis

We took the standard deviations $\sigma_{ERK}$ and $\sigma_A$ as a proxy for the amplitude in each signal. In simulations, we could simply average the amplitude - or amplitude ratio ($\sigma_A/\sigma_{ERK}$) - over all cells (Fig. 2B, S1A and S3C-F).

To acquire equivalent measurements for experiments we had to perform additional filtering and normalization steps, which we applied separately for each experimental repeat (N=3). Firstly, we removed cell tracks lasting less that 6 times the measured period of oscillation (period P=110 mins) and discarded cells with area measurement greater than 2.5 times the median value - which may occur for cells at the boundary of the Voronoi tessellation or wherever there are gaps in the cell detection due to weak ERK signal. We then interpolated signals closing across a maximum of two consecutive time-points (closing 15 minute gaps). After these steps we still retained large numbers of cell tracks (1252, 947, and 1083 for each repeat). We then non-dimensionalized area and ERK measurements by the mean across all cells so that units were equivalent to simulation. Finally, we applied filtered to remove transients longer than the period of ERK-density waves, e.g. from proliferation driven density changes, by subtracting a running mean with a window size of one period (110 mins).

From these processed signals, we calculated $\sigma_A/\sigma_{ERK}$ for each cell and plotted the average distribution across the three repeats (Fig. S3B). A log-normal distribution provided a good fit to this distribution, giving an estimate $\sigma_A/\sigma_{ERK} = 0.718 \pm 0.002$ with standard error estimated across the different repeats. To estimate $\sigma_A = 0.988 \pm 0.003$ and $\sigma_{ERK} \pm 0.113 \pm 0.004$ (mean and std - used to plot gray bands in Fig. S3D,E) we simply took the mean and standard deviation of the mean across the 3 experimental repeats.

### ERK auto-correlation

We used ERK auto-correlations (Fig. 3F and S2A-D) to study the decay in auto-correlation for different levels of noisy self-propulsion (Fig. 3G,H and S5B). Before calculating we first interpolated experimental ERK signals, closing up to two consecutive time-points (closing 15 minute gaps). We then cropped all time-courses to the same length as simulations (10 periods with $P = 110$ min for experimental and $P = 71$ min for simulation) and centered by subtracting the mean. Transient effects were less apparent in ERK signals (see insets Fig. S2D) compared to area so we did not apply further processing. After these steps we still had large numbers of cells: 939, 679, and 682 from each repeat.

We calculated the auto-correlation independently for each cell in both simulation and experiment:

$$\rho_{XX}(\tau) = E[X_{t+\tau}\bar{X}_t]/\sigma^2 \tag{1}$$

normalized by the variance so that $\rho_{XX}(0) = 1$.

To produce plots in Fig. S2B-D, we computed auto-correlation functions up to $\tau_{\max} = 3$ periods, comparing slices from time-courses $X(t)$ with a fixed length of 7 periods (total of 10 periods - $\tau_{\max}$). To study the decay in the first maximum, we only required to search up to $\tau = 1.5$ periods so we used a smaller $\tau_{\max}$ of two periods, slicing time-courses using a fixed window size of 8 periods. In order to show the slower decay in Fig. S2A we used $\tau_{\max} = 1,620$ with time-courses running for a total $T \approx 71,000$ min.

#### Amplitude of first maximum in ERK auto-correlation

We used the decay in amplitude of the first maximum in ERK auto-correlation to study the effect of noisy self-propulsion and to compare to experimental data. Given the stochasticity in individual cell auto-correlation functions (see Fig. S2D), we detected maxima only within a period $\tau$ of 1.5 times the expected period of oscillation, i.e. $P = 110$ min for data taken from the average auto-correlation over all cells (Fig. S2C) and $P \approx 71$ min for simulation as predicted from linear instability (see [1] and Fig. S1C). We show the results from this analysis in Fig. 3G,F and S5B.

#### ERK phase calculation and defect detection

We analysed the phase of ERK waves (definition Fig. S7 and see [5]) for two purposes: i) to quantify the effect of noisy self-propulsion ($f_0 > 0$) on the dynamics of ERK waves by comparing the average lifetime of phase defects (Fig. 3D,E and S5A) ii) to explore the effect of ERK-density waves on tissue fluidization (Fig. 4). In order to detect defects at the length scale of mechanochemical patterning, we first needed to smooth out the noise that self-propulsion introduced at the level of single cells. We also applied this procedure before the analysis of T1 transition locations to remove the effects of noise on our distributions. Below we explain our procedure for smoothing and defect detection.

Our first step was to create a 3D pixelized array of ERK values from simulation output or, for experiment, the output from cell tracking. For experimental data we also dropped the first 10 time-points to remove any initial transient. To pixelize each frame we used cell center positions to create a Voronoi tessellation and mapped ERK values to Voronoi cells, approximating each cell with an average of 10 pixels. We normalized ERK time-courses by centering and standardized each pixel in order to remove heterogeneity (experimental or biological) attributed to measuring ERK across different cells. Applying this normalization after pixelization meant that it was not necessary to have cells tracked frame-to-frame.

We then applied spatial smoothing to each frame using a Gaussian filter with a standard deviation of two cell widths, well below the expected wavelength of ERK-density waves ($\approx$ 20 cells). For simulations, we could wrap the filter across periodic boundaries, for experiment the input was extended by repeating values at the boundaries. Before calculating the ERK phase, we also applied a temporal band-pass filter to each pixel to recenter each time-course as well as to remove slow transients and high-frequency noise either side of the expected period of ERK oscillation or dominant mode. Specifically we filtered frequencies below $1/3 \times f$ and above $2.5 \times f$ where for experiment, $f$ (120 min) was chosen to be close to the dominant frequency measured from auto-correlation/Fourier analysis (Fig. 2C), and for simulations $f$ was estimated independently for each simulation from the average temporal Fourier spectrum of the spatially smoothed ERK signal. The above filtering was applied to the ERK signal rather than to the phase $\phi$ directly to avoid the computational cost of calculating circular means.

After smoothing, we calculated the ERK phase using the definition in Fig. S7 (see also [5]) with $\tau$ set to approximately 1/4 the measured period: 30 min for experiment or taken from the independent spectral analysis of the dominant mode for each simulation. Finally, we detected defects by integrating anti-clockwise around overlapping 2x2 pixel squares, assigning charge $\pm 1$ for integral values $\pm 2\pi$.

#### Identification of T1 transition events

The location of T1 transition sites was output automatically by the Chaste vertex model simulation software [3]. Inspection of this output revealed that edges would occasionally swap back and forth rapidly between time-steps before a T1 was completed or rejected, leading to an over-count of T1 events. After correcting for this we validated the location of T1 events by overlaying them (circles) on vertex diagrams across consecutive frames (Fig. 4A).

#### ERK phase at T1 location distributions

We created phase distributions by mapping ERK phase to the position of T1 transitions (Fig. 4B). After noticing asymmetry in the results of our initial analysis (Fig. S8 - green) we applied two different corrections to remove possible bias: one aimed to correct for the non-uniform distribution of total phase computed for each simulation across all of space and time (blue histogram Fig. S8) which might stem from non-linear temporal oscillations or a miss-match between $\tau$ used to calculate the phase and the true period of oscillation, i.e. if the cells spend more time at a certain $\phi$ then there is a bias towards detecting more T1s with this value.

Applying this first correction presents a second issue: ERK is associated to cells of differing area and the distribution of $\phi$ should indeed be non-uniform because of the correlation between ERK and cell size. We therefore applied a second correction, weighting each T1 event by the local cell area to account for the higher number of cell junctions at higher cell density. For the area correction, we also have to consider that there is a short delay between area and ERK. We explored offsetting the area weighting by one or two time-steps but did not observe any substantial difference. In the final corrected distributions (Fig. S8 - black and Fig. 4B) the level of asymmetry is much reduced.

#### Defect tracking and defect lifetimes

We used the python package trackpy [6] to track ERK phase defects, both in simulations and in experiments. We took advantage of the $\pm$ sign to track defects of opposite charge separately before recombining results. For experimental data, we used a search range of 2 cell widths for the maximum displacement between frames and allowed gap closing for a maximum of one missing frame. For simulations we used a search range of 1.5 cell widths for the maximum displacement between frames with the same gap closing. The slightly different parameters were chosen on account of the different time gap between frames (5 min for experiments vs 3 min for simulations) and based on manual calibration and validation from visual inspection of the tracking output (see Movies S2-4).

Before plotting defect lifetime distributions, we applied several steps to remove bias. In experiments, we excluded tracks passing within $\approx 3$ cell widths of a boundary to exclude defects that exit the field of view. For simulations, we were able to track defects across the periodic boundaries. We also discarded tracks that were present at the start or end time-points as these were not fully observed. Finally, we weighted lifetimes $\tau$ by $1/(T-\tau)$, where $T$ is the total time-span of the simulation or experiment to account for the changing size of observation window in which a defect could be born and fully observed until death. We then used these weighted lifetimes to construct histograms using fixed bin edges - 10 bins of 50 minute widths from 0-500 minutes.

We observed that, apart from the first bin, the lifetime distributions were well approximated by an exponential distribution (see e.g. Fig. 3D and similar observations in [5]). Since remaining noise in ERK signal or bias from tracking errors might contribute towards a greater than expected number of short-lived defects populating the first bin, we reasoned that we would obtain truer measurements of average defect lifetime using parametric estimates from fits of an exponential distribution in which the first bin was excluded.

For fitting experimental distributions, we used the mean and variance of each bin across three repeats to perform a weighted least squares regression in semilog-space (Fig. 3D), giving an estimate $\tau = 79 \pm 6$ min including standard error calculated from individual unweighted fits of the three repeats. For simulations we instead fit individually using ordinary least squares and averaged the results across repeats initialized from different random seeds to produce the plots in Figs. 3E and S5A.

### MSD analysis

We analysed mean square cell displacements to study how increasing self-propulsion strength $f_0$ affects tissue fluidity in the presence of mechanochemical waves (Fig. S6B-D). For simulations, we could simply calculate mean square cell displacement from the start of simulation output (after burn-in) and average across all cells. For experiments, we took the data from cell tracking and retained only tracks longer 100 frames (500 minutes), leaving 2827, 2529 and 3193 cell tracks in each repeat. We then calculated the mean square displacement from initial position for each cell vs time before averaging values over cells in each repeat. We repeated this analysis while removing the average displacement of the entire tissue to correct for global flows or movement of the sample. In Fig. S6D we plot both versions for the average across three repeats.

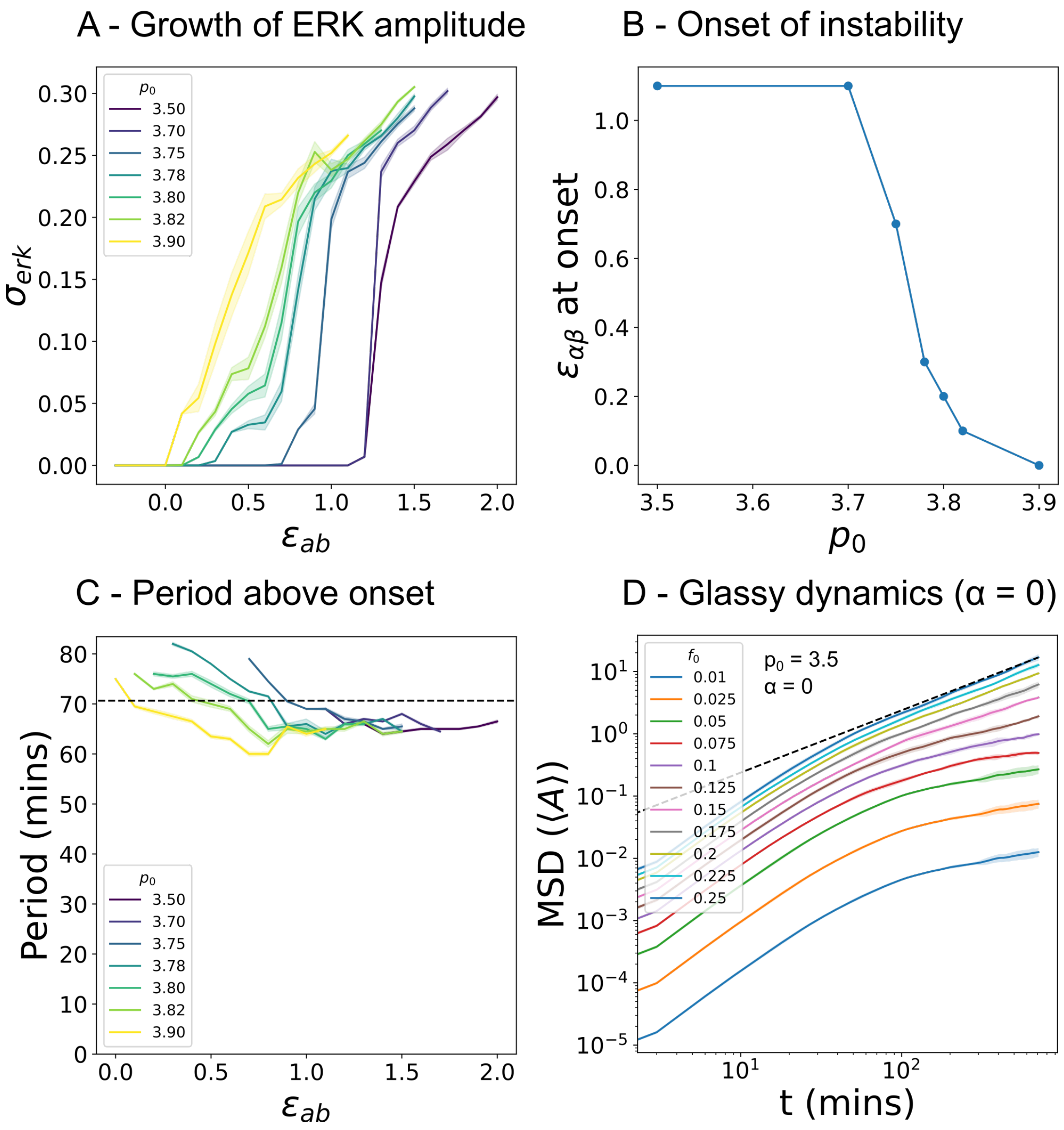


FIG. 1: Quantitative Phase diagram of ERK patterning and glassy dynamics as a function of system parameters. A/ Amplitude of ERK oscillation vs normalized mechanochemical coupling strength $\epsilon_{\alpha\beta} = (\alpha\beta - \alpha\beta_c)/\alpha\beta_c$ in simulations of the 2D vertex model where $\alpha\beta_c$ is the value for the onset of instability predicted by the 1D linear instability analysis in [1]. Colors indicate different values of the shape index $p_0$ with solid-fluid transition expected at critical value $p_0^* \approx 3.81$ [7]. Errors show standard error calculated over 6 repeats starting from different initial seeds. B/ Values of the critical value of mechanochemical coupling strength $\epsilon_{\alpha\beta}$ at onset of instability vs the shape index $p_0$ corresponding to the plots shown in A. C/ Temporal period of ERK oscillations above the instability onset as measured from location of the first maximum in ERK temporal auto-correlation function. The dashed line indicates the theoretical prediction from [1]. Error bars indicate standard error over the same simulation repeats as in A. D/ Average cell mean square displacement (MSD) vs time for $p_0 = 3.5$ and different values of self-propulsion force $f_0$ in the absence of mechanochemical ERK waves ($\alpha = 0$). Errorbars indicate standard deviations over 6 repeats from different initial seeds. The dashed black line indicates the linear scaling MSD $\sim t$ expected in the diffusive limit, while sub-diffusive scales are expected for caged motion below a critical self-propulsion force. Recovery of the diffusive limit with increasing $f_0$ indicates a glassy rheology with noisy self-propulsion fluidizing the otherwise solid tissue.

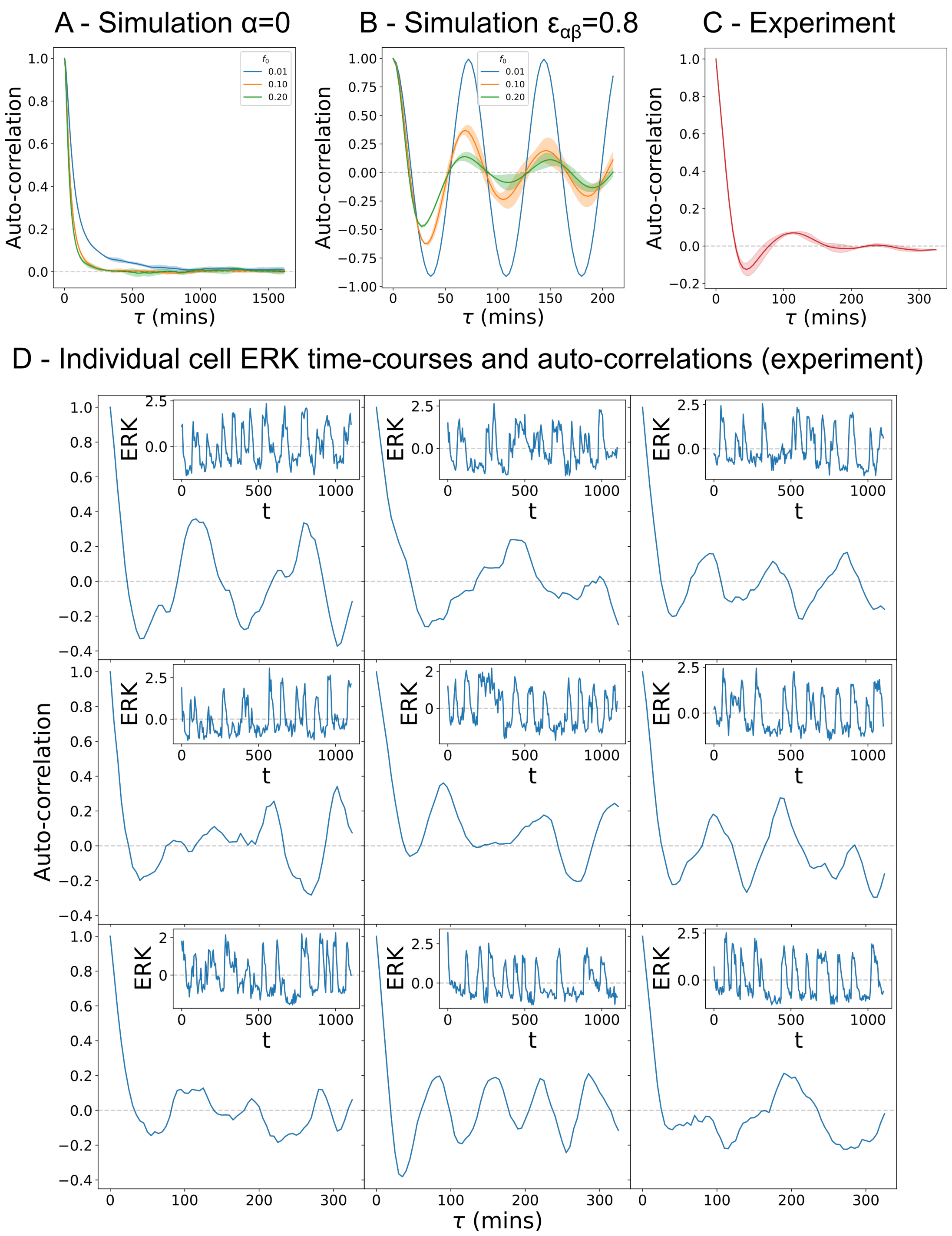


FIG. 2: Quantifications of ERK dynamics in simulation and experiments. A,B/ Temporal auto-correlation in ERK signal averaged over $44 \times 44$ cells from vertex model simulations with varying strengths of self-propulsion force $f_0$, with and without the presence of ERK-density oscillatory waves: $\alpha\beta = 0$ and $\epsilon_{\alpha\beta} = 0.8$. Errorbars show respectively standard deviations over 6 and 5 simulations initialized from different seeds. C/ The equivalent average cell auto-correlation calculated for ERK data from MCDK monolayers with error bars indicating standard deviation over N=3 experimental repeats from [1]. D/ ERK timecourses and corresponding autocorrelations for individual cells randomly selected across 3 experimental repeats (each column a separate repeat).

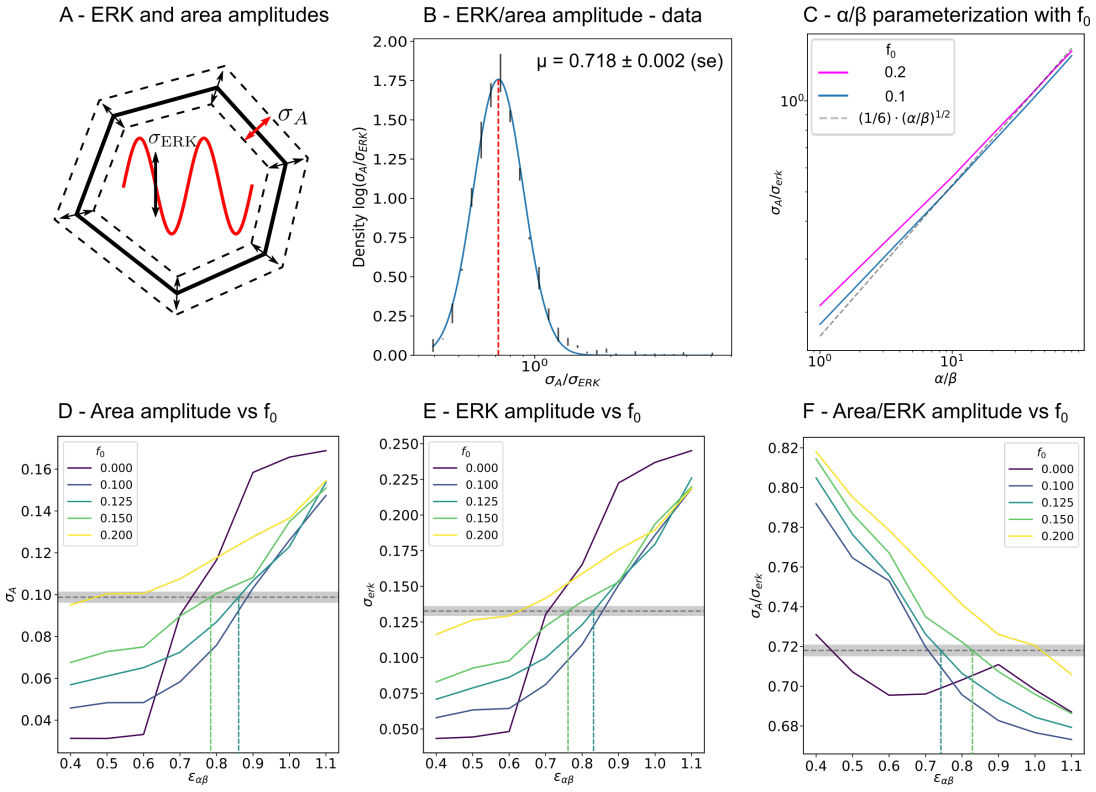


FIG. 3: Parameter fitting strategy. A/ Schematic of the temporal area and ERK signal amplitudes used in our analysis. The relative amplitude of $\sigma_A/\sigma_E$ scales with the ratio $\alpha/\beta$ (see Fig. 1B) in our model. B/ Average experimental distribution of area to ERK amplitude ratio with lines to indicate mean and and log-normal fit. Errorbars indicate standard deviation over N=3 repeats from [1]. C/ Repeat of the analysis in Fig. 2B (grey dashes) for simulations introducing active self-propulsion, showing the robustness of our estimate for $\alpha/\beta$ with respect to different values self-propulsion force $f_0 > 0$. D-F/ Absolute area and ERK amplitude and area to ERK amplitude ratio vs strength of mechanochemical coupling for different values of $f_0$. Dashed line shows estimated means and standard deviations ($\sigma_A = 0.988 \pm 0.003$, $\sigma_{ERK} = 0.133 \pm 0.004$, $\sigma_A/\sigma_{ERK} = 0.718 \pm 0.003$) from N=3 experimental repeats, allowing us to identify a compatible region in parameter space of $\epsilon_{\alpha\beta} \approx 0.8$ and $f_0 \sim 0.125 - 0.15$.

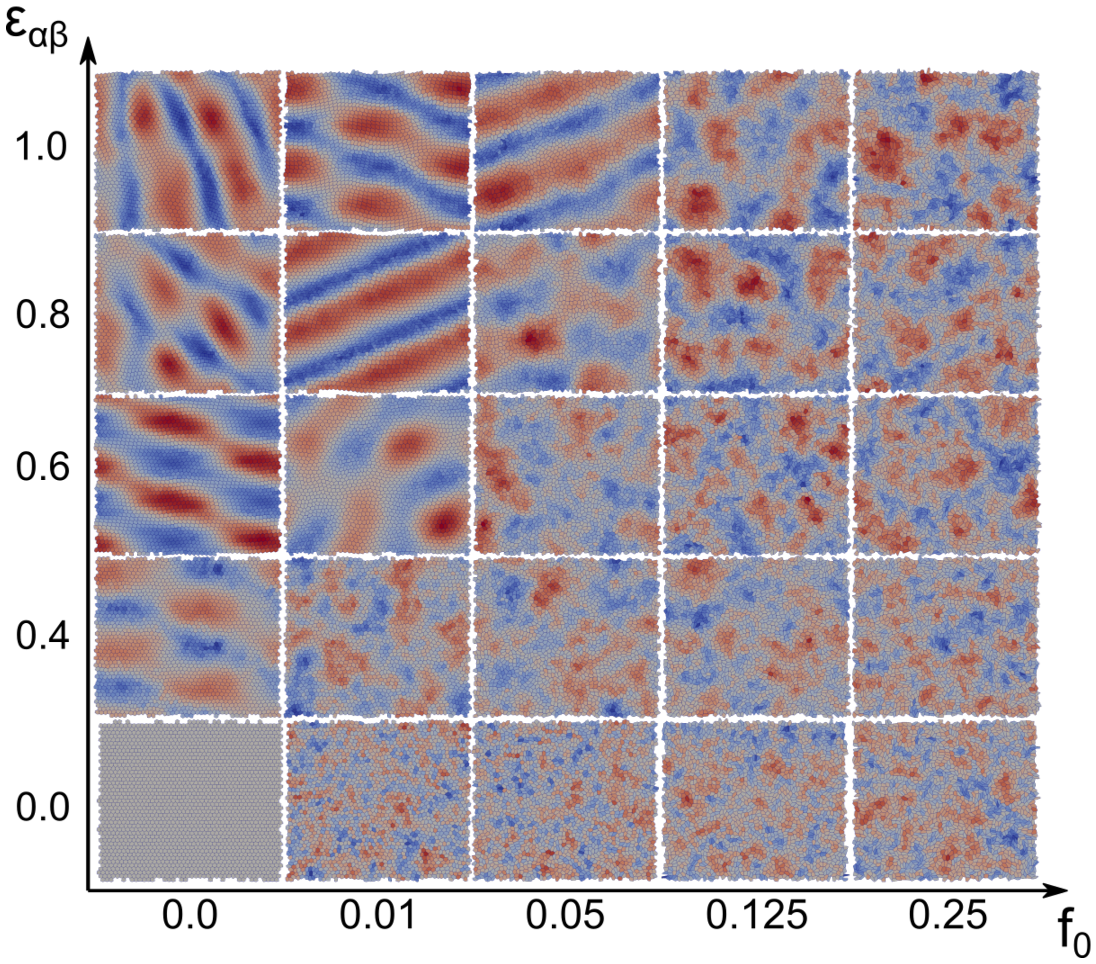


FIG. 4: Phase diagram of monolayer dynamics as a function of self-propulsion force $f_0$ and the absolute strength mechanical of chemical coupling $\epsilon_{\alpha\beta} = (\alpha\beta - \alpha\beta_{\rm c})/\alpha\beta_{\rm c}$. Fig. 2C showed a similar phase diagram as a function instead of the ratio $\alpha/\beta$.

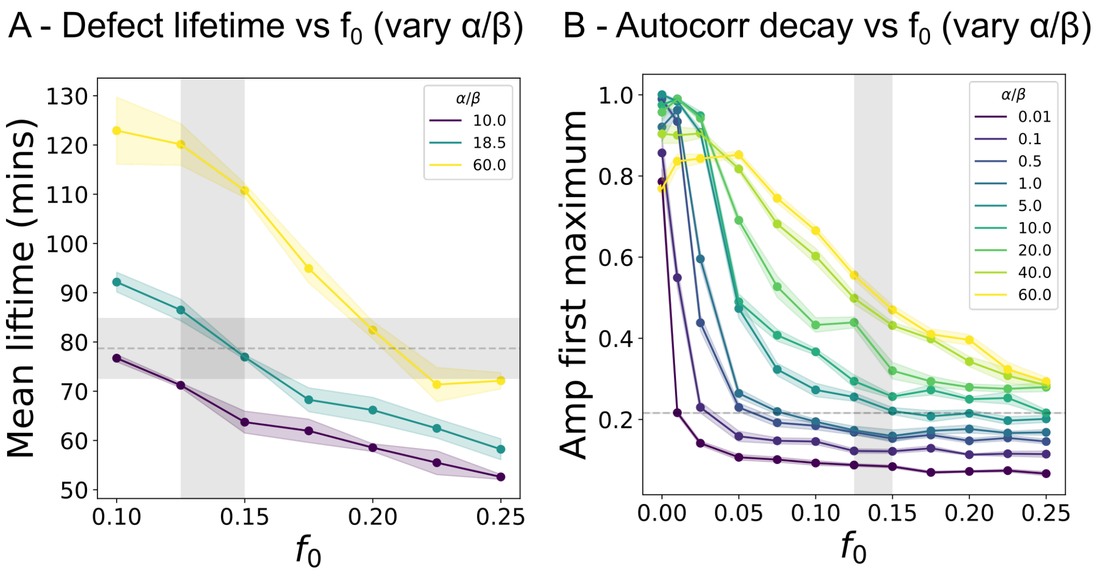


FIG. 5: Additional comparisons between model prediction and experimental data. A,B/ Mean defect lifetime and amplitude of first maximum in ERK auto-correlation distributions (see also Fig. 3E,H), as the ratio of mechanochemical coupling $\alpha/\beta$ is varied in simulations. Errorbars on curves indicate standard errors from respectively 3 and 6 simulation repeats initialized from different seeds. Along the y-axes grey shaded regions indicate measurements from defect lifetimes and ERK temporal autocorrelations in data (Fig. 2D,G) and along the x-axes the parameter region for $f_0$ inferred from the analysis of area and ERK signal amplitudes (Fig. S3D-F). All simulations used best estimate $\epsilon_{\alpha\beta} = 0.8$ from the same analysis of amplitudes (Fig. 3D-F).

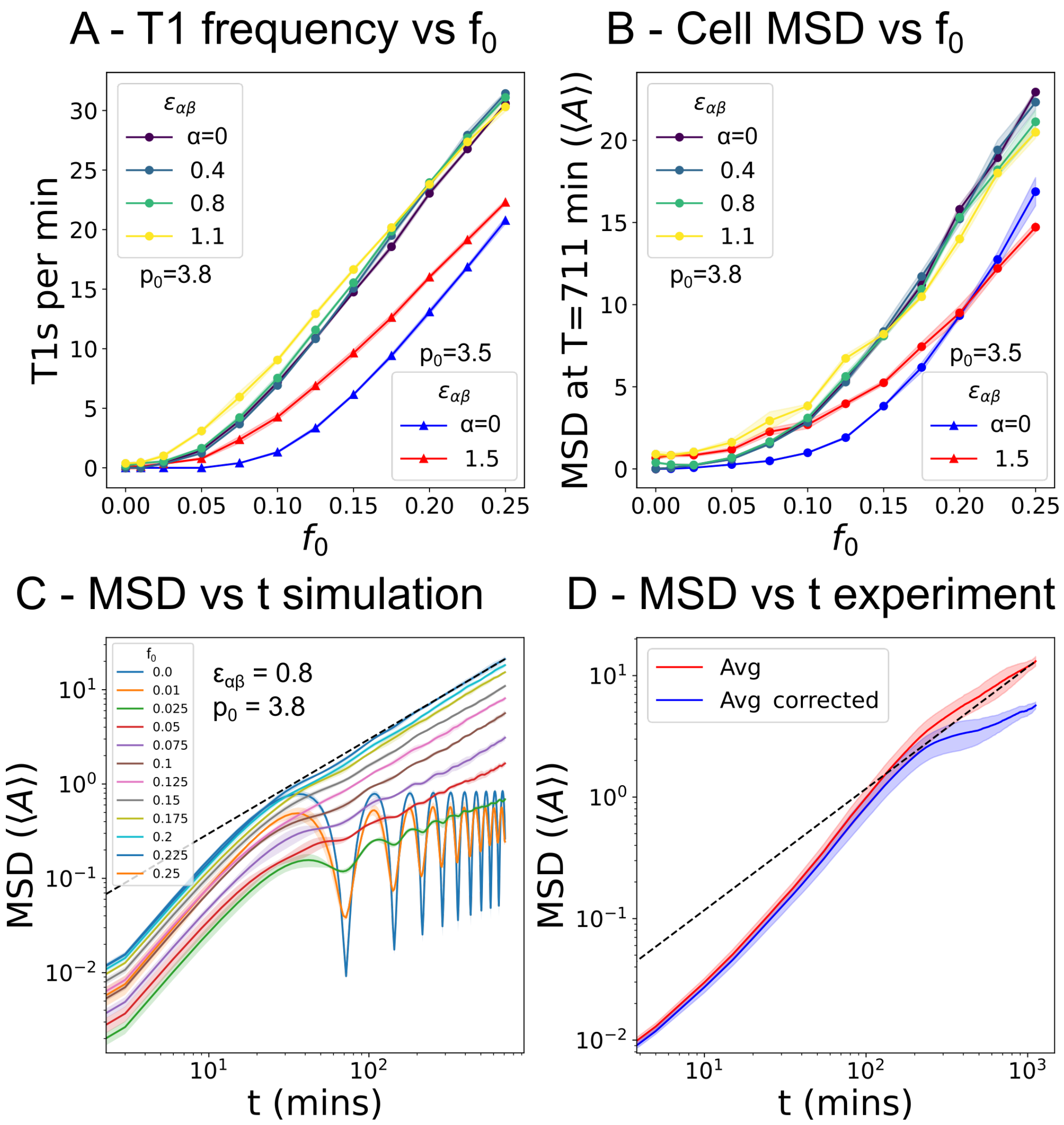


FIG. 6: Impact of mechanochemical couplings on local tissue fluidity and dynamics. A,B/ Cell-cell rearrangement, or T1 transition, frequency (A) and cell mean square displacement (B) (in units of average cell area) vs self-propulsion force $f_0$ for different values of mechanochemical coupling strength $\epsilon_{\alpha\beta}$ and shape index $p_0$. Errorbars indicate standard errors over 4 repeats ($p_0 = 3.8$) and 6 repeats ($p_0 = 3.5$). C/ Average cell mean square displacement vs time in simulations for different values different values of $\epsilon_{\alpha\beta}$ and $f_0$. Errorbars indicate standard deviations over 4 repeats. D/ Average cell mean square displacement vs time over three separate experimental repeats. Errorbars indicate standard deviation before (red) and after (blue) correcting for the average global translation of cells. Dashed black lines in C and D indicate the linear scaling MSD $\sim t$ expected in the diffusive limit.

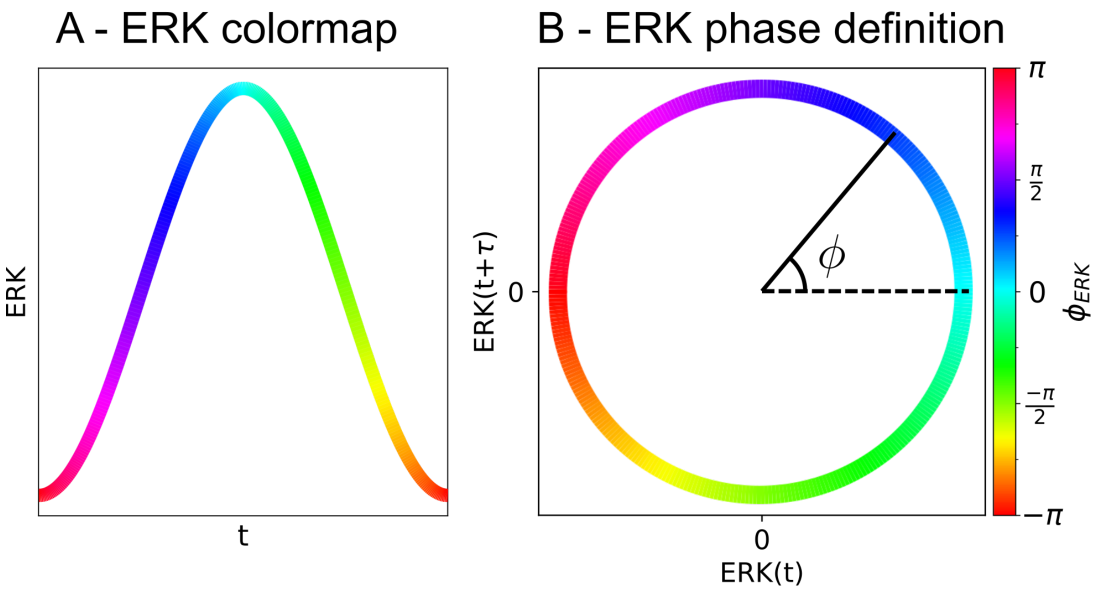


FIG. 7: Schematic of the color scheme used in the main figures. A/ Colormap of ERK phase $\phi$ overlayed on an illustrative ERK oscillation cycle. B/ Definition of ERK phase $\phi$ using the cycle formed between ERK signal and ERK signal with a delay $\tau$ of one quarter of a period of oscillation (see [5]).

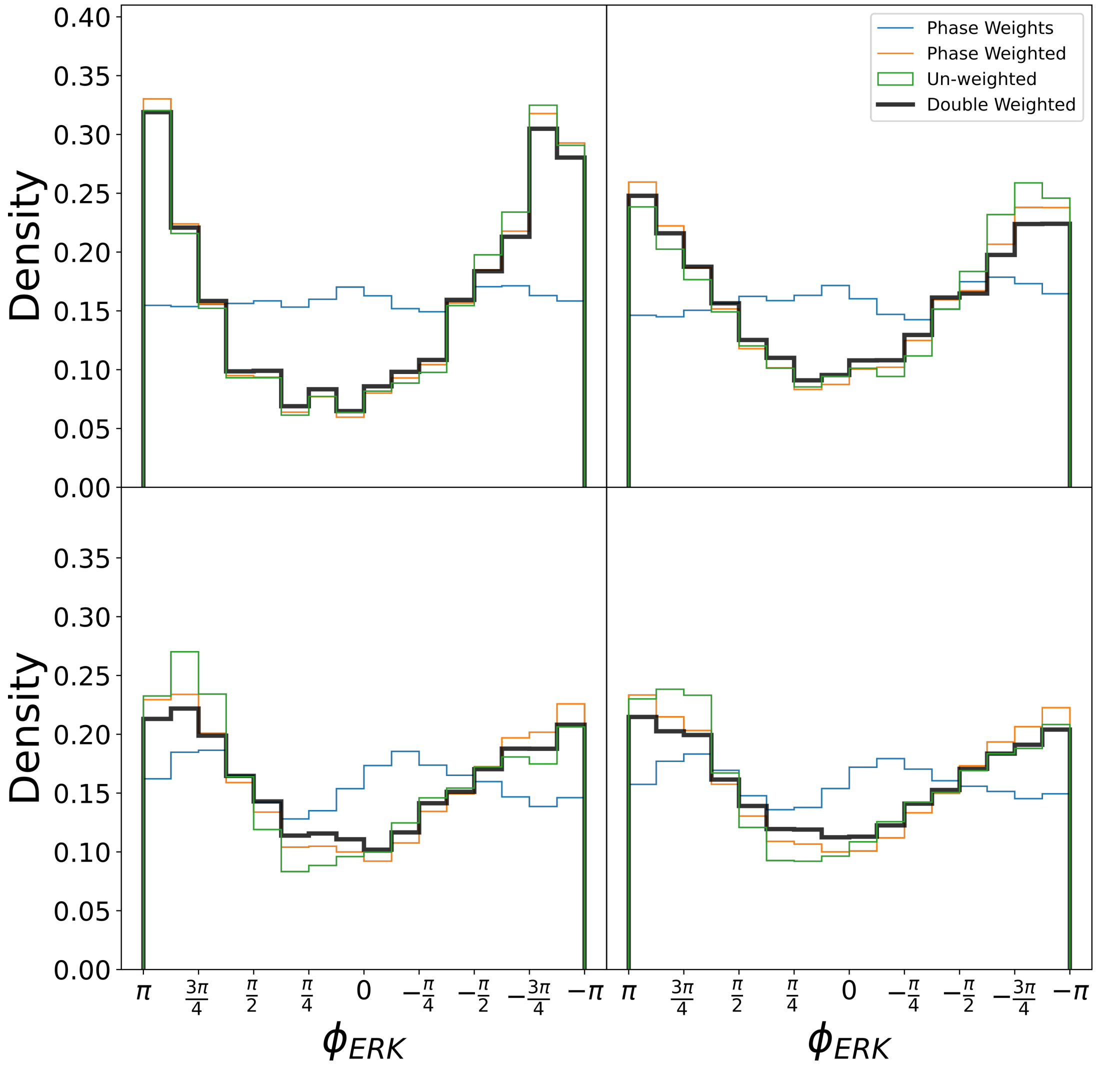


FIG. 8: Illustration of different corrections applied to the distribution of ERK phase at T1 transition sites. Black lines correspond to the histograms presented in Fig. 4B. Corrections for non-uniform phase distributions (blue) and local vertex density removed much of the asymmetry present in the unweighted histograms (green). See methods section "ERK phase at T1 location distributions" for further details.

TABLE I: Vertex model parameters: Time-scales $\tau_E$ and $\tau_A$ from [1], $\tau_p$ consistent with traction force measurements from [1, 8]. Ratio $K_A/K_p$ extracted from [9] (same as used in [10]) with substrate friction re-scaled to give a wavelength of instability of around 20. Non-dimensionalization used average cell area estimated from confluent data and timescale of ERK activation $\tau_E$. See methods section "Vertex model simulations" for further details.

| Parameter | Value | units | Non-dimensionalized |
|---|---|---|---|
| $K_A/\mu$ | 0.032 | $h^{-1} \cdot \mu m^{-2}$ | 1.2 |
| $K_P/\mu$ | 3.8 | $h^{-1}$ | 0.38 |
| $\tau_E$ | 0.1 | $h$ | 1 |
| $\tau_A$ | 2 | $h$ | 20 |
| $\tau_p$ | 0.5 | $h$ | 5 |
| Avg Cell Area | 380 | $\mu m^2$ | 1 |

### Movies

- Movie S1. Vertex model simulation for best fit parameters $\alpha/\beta = 18.5$, $\epsilon_{\alpha\beta} = 1.8$ and $f_0 = 0.15$ with colors to indicate levels of ERK activation.
- Movie S2. ERK phase and defect tracking for simulation with lower $f_0 = 0.10$. Other parameters $\alpha/\beta = 18.5$ and $\epsilon_{\alpha\beta} = 1.8$.
- Movie S3. ERK phase and defect tracking for simulation with higher $f_0 = 0.25$. Other parameters $\alpha/\beta = 18.5$ and $\epsilon_{\alpha\beta} = 1.8$.
- Movie S4. ERK phase and defect tracking for one the three experimental repeats.
